# Foster thy young: Enhanced prediction of orphan genes in assembled genomes

**DOI:** 10.1101/2019.12.17.880294

**Authors:** Jing Li, Urminder Singh, Priyanka Bhandary, Jacqueline Campbell, Zebulun Arendsee, Arun S. Seetharam, Eve Syrkin Wurtele

**Author notes:** Equal first authors (Jing Li, Urminder Singh).

## Abstract

Proteins encoded by newly-emerged genes (“orphan genes”) share no sequence similarity with proteins in any other species. They provide organisms with a reservoir of genetic elements to quickly respond to changing selection pressures. Here, we systematically assess the ability of five gene annotation pipelines to accurately predict genes in genomes according to phylostratal origin. BRAKER and MAKER are existing, popular *ab initio* tools that infer gene structures by machine learning. Direct Inference is an evidence-based pipeline we developed to predict gene structures from alignments of RNA-Seq data. The BIND pipeline integrates *ab initio* predictions of BRAKER and Direct inference; MIND combines Direct Inference and MAKER predictions. We use highly-curated Arabidopsis and yeast annotations as gold-standard benchmarks, and cross-validate in rice. Each pipeline under-predicts orphan genes (as few as 11 percent, under one prediction scenario). Increasing RNA-Seq diversity greatly improves prediction efficacy. The combined methods (BIND and MIND) yield best predictions overall, BIND identifying 68% of annotated orphan genes and 99% of ancient genes in Arabidopsis. We provide a light weight, flexible, reproducible solution to improve gene prediction.

## MAIN

Eukaryotic and prokaryotic genomes contain genes (“orphan genes”) whose proteins are recognizable only in a single species. Some of these have emerged *de novo* from the genome, while others have diverged from their cousins so quickly that no sequence similarity is detectable^1–5^.

As encoders of completely novel proteins, orphan genes provide a disruptive force in evolution. Orphans play a crucial role in adaptation to new biological niches. Studies from vertebrates, annelids, insects, fungi, and plants show that many extant orphan proteins mitigate novel biotic challenges (prey, predators, hosts) or emergent environmental shifts^2,6–9^.

Some proteins encoded by orphan genes (e.g., toxins^6^) act externally, while others integrate into internal metabolic and developmental pathways^7^. Phylostratigraphic reconstructions have implicated orphans in the evolution of new reproductive and neural structures^10,11^. Thus, the advent of orphan genes may provide a critical enabler of speciation. The ability to accurately predict orphan genes and other young lineage-specific genes conveys unique percipience about evolution and ecology^2,11,12^.

A subset of orphans are retained as genes and continue to evolve, such that each genome contains a mixture of genes of different ages (phylostrata)^1,3,13,14^. Thus, the age of each gene can be considered the time since its deepest ancestor emerged (**Figure 1**), as opposed to its most recent duplication. Even the most ancient genes were orphan genes once (for protein-coding genes, these would be genes whose proteins trace back to early eukaryotic organisms or to prokaryotes). Of the estimated billions of extant protein-coding orphan genes across all eukaryotic species (conservatively calculated as 8.7 million extant eukaryotic species^15^ × 1000 orphans per eukaryote)^16^, the functions of only a few hundred have been elucidated^2,3,7,8,17–21^. In contrast, about two thirds of the annotated genes of Arabidopsis are very ancient^2^, tracing back to a very early eukaryotic or a prokaryotic origin. In part because they have common motifs across species, these ancient genes are more likely to be at least partially characterized.

**Figure 1.**
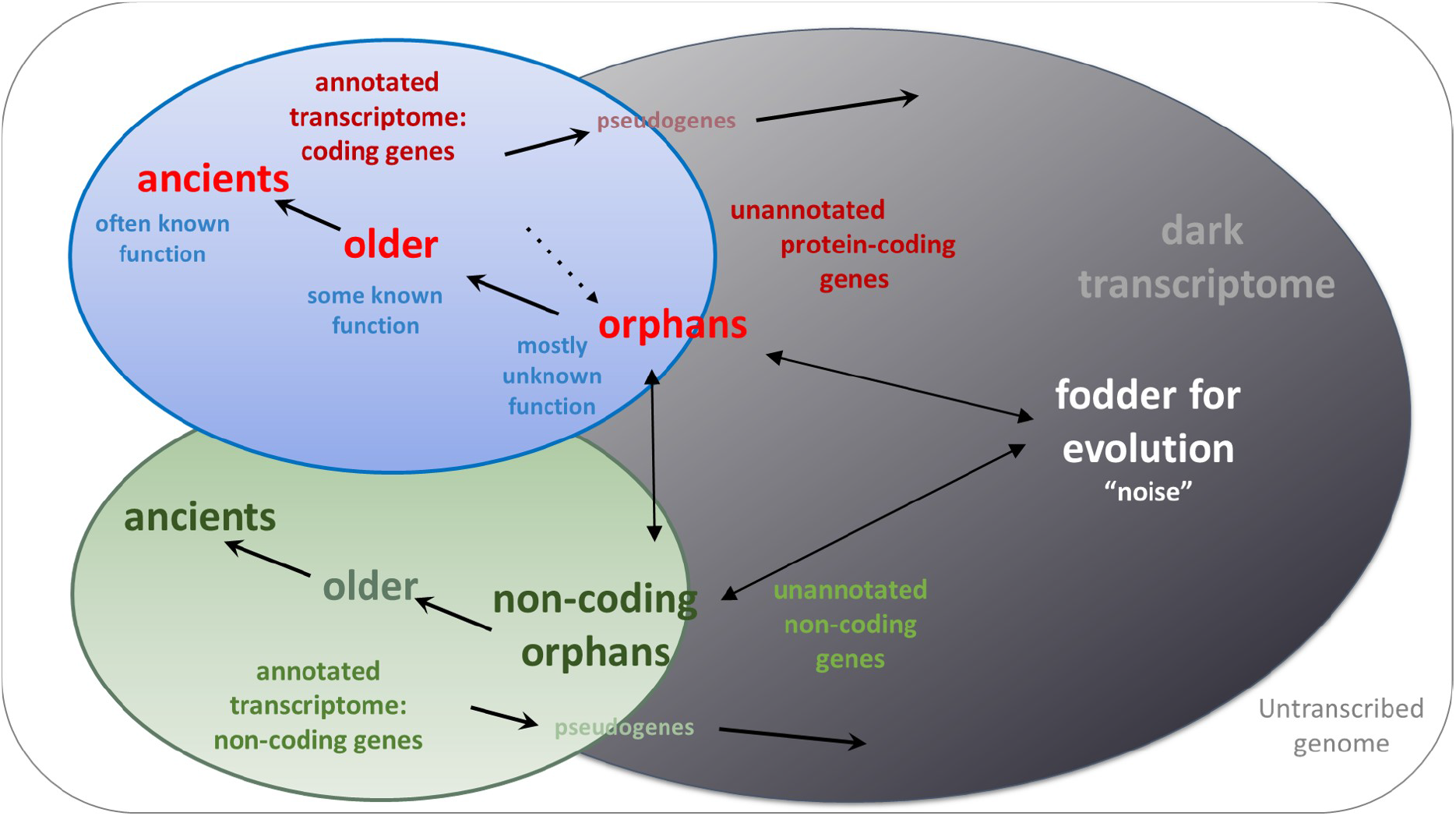
The dynamic transcriptome. Genes form, evolve, remodel, and recede in a continuum across evolutionary time. These dynamics result in a significant challenge to gene annotation. The expressed genome, i.e., the transcriptome, is comprised in part by annotated genes. However, the transcriptome also includes a vast, uncharacterized but expressed body of sequences that can be termed the “*dark transcriptome*”. Within this dark transcriptome is low abundance non-genic “noise”, as well as (as-yet-unannotated) protein-coding genes, pseudogenes, and non-coding genes^5,26,48,50–52^. Protein coding orphan genes can be formed from expressed non-functional sequence, extant non-coding genes, and to a lesser extent from existing genes whose proteins have rapidly evolved beyond recognition^4,12,27^. Over evolutionary time, some orphans will be retained in the species and become members of deeper phylostrata. In general, an older, more conserved gene is more likely to have a known molecular function. The proportion of functional genes that are unannotated in any given species is unclear; we posit that, depending on the species, a sizable proportion of orphan genes remain unannotated. This is because many are sparsely expressed^16,25,44,48^, by definition none have homologs, many may have not yet evolved the canonical features by which a gene can be recognized *ab initio*, and there is a grey area in evolution between “noise” and “gene”. Black arrows, evolutionary transitions; red font, protein-coding genes; green font, non-[protein]-coding genes; grey font, non-genic transcripts; blue oval, annotated protein-coding genes; green oval, annotated non-[protein]-coding genes.

The technique of phylostratigraphy identifies genes based on the earliest common ancestor to which a protein homolog can be traced^1^. Gene “hybrids”, incorporating domains of several phylostratal origins, are assigned according to their most ancient domain. Phylostratigraphy classifies genes independently of their origin, including those that emerged *de novo* from non-genic sequence or from within an existing gene, and those that evolved so rapidly that they cannot be recognized even in genomes of their closest (genus-level) relatives^1–3^. Phylostratigraphy has been rendered reproducible and automated via *phylostratr* software^13^.

Gene annotation is a fundamental step in genome sequencing projects, however, no standard best practice has been established, and protocols are often diverse and not well-documented (**Supplementary Table 1**). Prevailing methods often combine *homology-based analysis*, which compares a new genome to previously-identified genes from other species, and *ab initio prediction* of genes from the genome sequence^22^. Each approach may have inherent bias against orphan genes^23^. Homology-based methods assume that genes have identifiable orthologs in other species. Because orphan genes are species-specific, homology-based methods are not useful in predicting them. *Ab initio*-based predictors assume a pre-defined gene structure for all protein-coding genes of an organism^24^.

Nucleic acid signatures by which genes are predicted *ab initio* can include sequence motifs of untranslated regions, translation start sites, termination sites, and intron-exon boundaries. However, canonical sequence signatures may be less well-defined in young genes^25^. in which case *ab initio* approaches might be less likely to detect them. The ability of *ab initio* approaches to detect genes of recent origin had not been directly evaluated.

A more straightforward evidence-based approach to identify genes is to directly align RNA-Seq data to genomes^26,27^. This approach is key for predicting non-coding RNAs^28^ and young genes^3^.However, it has been less widely adopted to annotate protein coding genes, in part because of the challenge of distinguishing “noise” from true genetic signal^26,27^. One approach to reduce “noise” and other false positive predictions is to combine direct inference of genes with sequence similarity^24,26^; this approach excludes orphan genes^29^.

Here, we compare predictions of five annotation pipelines: MAKER^30^, BRAKER^31^, Direct Inference- an evidence-based pipeline we developed to predict genes by genome-guided alignment of RNA-Seq data, and two novel pipelines that combine *ab initio* and the Direct Inference approaches: **M**AKER-**In**ferred **D**irectly (MIND), and **B**RAKER-**In**ferred **D**irectly (BIND) (**Figure 2**).

**Figure 2.**
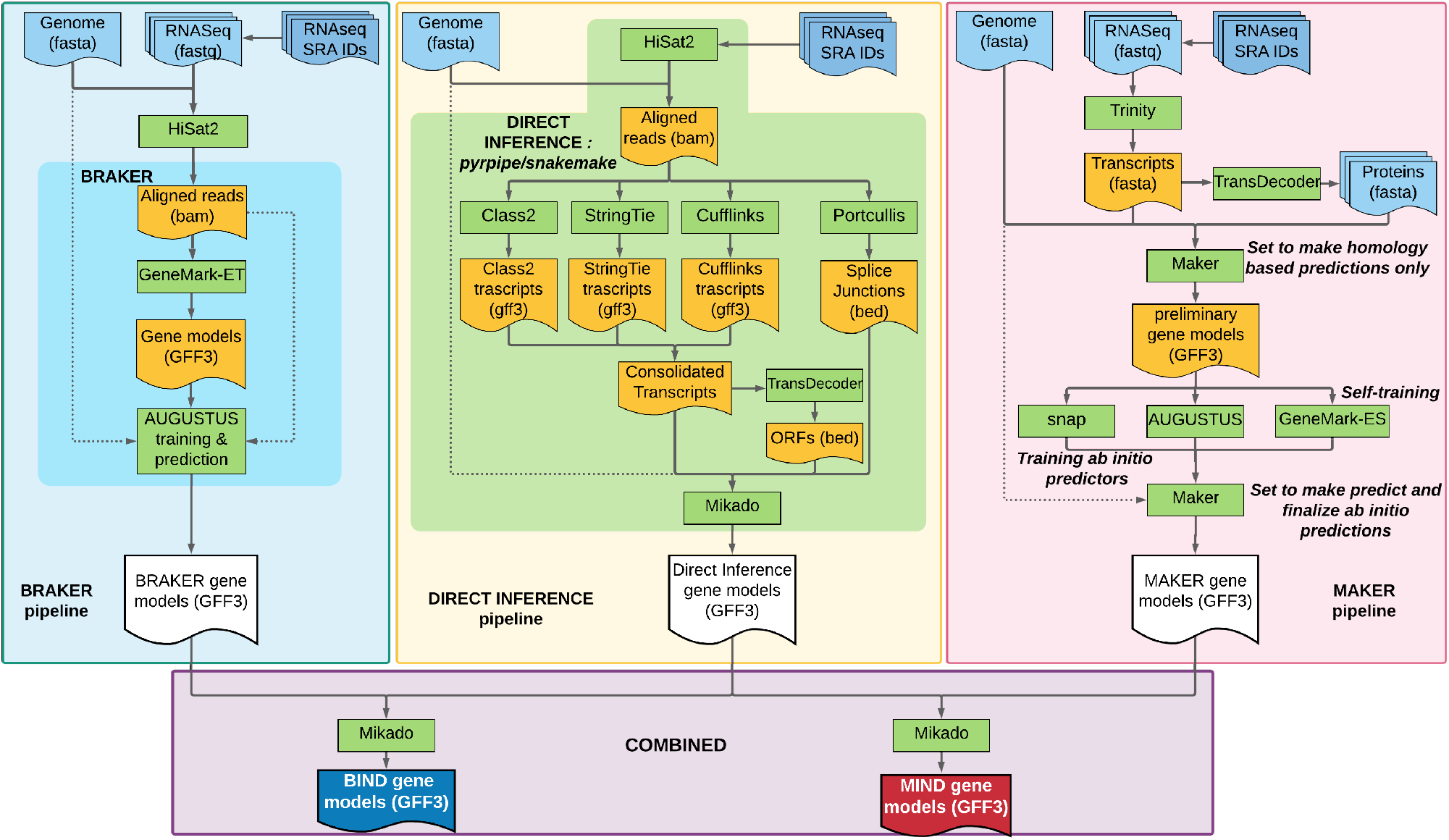
Schematic diagram of MIND and BIND gene prediction pipelines. BRAKER^31^ (top left) and MAKER^30^ (top right) predict genes based solely on *ab initio* machine learning, while Direct Inference (top middle) is an evidence-based method we developed to predict genes based on alignment of RNA-seq data to an assembled transcriptome. BRAKER (darker turquoise area) automates most of the pipeline software; Direct Inference (lime green area) automates the full pipeline; MAKER requires user-intervention for each step. BRAKER parameters for training are hard-coded within script and difficult to manipulate; Direct Inference and MAKER enable parameter management. Direct Inference also enables software to be exchanged (e.g., STAR for HiSat2)^34^, MAKER uses settings files and can be user-edited to change parameters. BRAKER, MAKER, and Direct Inference prerequisites and features are shown in **Supplementary Figure 7**. The *BIND* and *MIND* pipelines (bottom) use Mikado^77^ to combine predictions from the Direct Inference pipeline with predictions of either MAKER (*MIND*) or BRAKER (*BIND*). The full *MIND* and *BIND* pipelines with clearly documented, open source code are at https://github.com/eswlab/orphan-prediction.

We compare our gene predictions to those of the highly-curated “gold-standard” gene annotations in *Arabidopsis thaliana* (Arabidopsis) and *Saccharomyces cerevisia* (yeast) and apply these methods to the much newer and hence less-curated NCBI annotations for a genome of the staple crop, *Oryza sativa* (rice)^32^. Our results reveal that *ab initio* prediction pipelines can vastly under-detect younger genes. We show that diverse RNA-Seq evidence significantly improves gene annotation, in particular for younger genes. We demonstrate that the novel BIND and MIND pipelines improve the number and performance of predictions.

To enable Findable, Accessible, Interoperable and Reusable (FAIR)^33^ pipelines for MIND and BIND, we implemented the Direct Inference pipeline in an automated, reproducible manner using the python-based RNA-Seq processing workflow, pyrpipe^34^ such that it can be easily customized; we included singularity containers for MAKER and BRAKER.

## Results

MAKER is a popular software that combines a variety of *ab initio* approaches to predict genes^30^. Our original impetus for this research was to assess MAKER’s ability to predict orphan genes. However, our initial analysis predicted only 11% of the orphan genes annotated in The Arabidopsis Informatics Resource (TAIR; Araport11 version)^35^.

This poor performance of MAKER in identifying orphans and other young genes led us to consider other gene prediction scenarios. First, we reasoned that using more RNA-Seq evidence for training MAKER’s algorithms might improve predictions. However, there was little guidance in the literature on how to formulate the training datasets for *ab initio* gene predictions^30,31,36^; many genome annotations use very limited datasets (**See Supplementary Table 1**). Second, we considered that other gene prediction programs might improve prediction outcomes.

Thus, we evaluated the efficacy of different gene prediction software and the extrinsic evidence of varied sizes on predictions. Specifically, raw reads from diverse RNA-Seq datasets were used by *ab initio* (MAKER and BRAKER), Direct Inference or the combined (MIND and BIND) approaches to predict genes.

We focused specifically on predicting protein-coding genes; the pipelines we applied - MAKER, BRAKER, and Direct Inference-each require the presence of an ORF to indicate the potential for translation. Protein evidence for each ORF would confirm its translation.

### Arabidopsis gene predictions by MAKER

MAKER^30^ is a widely-used gene prediction pipeline. Reads are aligned to the genome and assembled into transcripts before being provided to MAKER, which uses assembled transcripts of RNA-Seq datasets along with predicted protein sequence exclusively as training data.

We assembled seven combinations of transcript and protein evidence (**Table 1 and Supplementary Tables 2-A and 3-A** and provided these to MAKER as training data. Gene predictions corresponding to annotated genes were greater when RNA-Seq and protein data were both provided **Supplementary Table 4-A**. Regardless of input evidence, MAKER’s ability to predict genes (**Figure 3** and **Supplementary Table 4-A**) was greatest for the genes of the oldest phylostratum (Cellular Organisms, PS1) and progressively decreased for younger phylostrata.

**Table 1.** Gene prediction scenario used for *A. thaliana*. The “Typical” RNA-Seq dataset is of a size similar to or greater than that used in many gene annotation projects. The “Pooled” dataset is more diverse and includes the Typical dataset. The “Orphan-rich” dataset is designed to maximize orphan representation.

| Scenario (abbreviation) | Analysis Pipeline | Description | Extrinsic Evidence |  |
| --- | --- | --- | --- | --- |
|  |  |  | # SRR samples | Datasize (GB) |
| Ara11 | NA (gold-standard annotations from TAIR<sup>35</sup>) | Araport11 version of *A. thaliana* annotations | NA | NA |
| Maker-Typical | MAKER | Maker predictions using Typical dataset (assembled transcripts and translated proteins) | 12 | 12.8 |
| Maker-Pool |  | Maker predictions using Pooled dataset (assembled transcripts and translated proteins) | 77 | 241.4 |
| Maker-Orphan |  | Maker predictions using Orphan-rich dataset (assembled transcripts and translated proteins) | 38 | 595.1 |
| Braker-Typical | BRAKER | Braker predictions using Typical RNA-Seq dataset (raw RNA-Seq) | 12 | 12.8 |
| Braker-Pool |  | Braker predictions using Pooled RNA-Seq dataset (raw RNA-Seq) | 77 | 241.4 |
| Braker-Orphan |  | Braker predictions using Orphan-rich RNA-Seq dataset (raw RNA-Seq) | 38 | 595.1 |
| DirInf-Typical | Direct Inference | Direct Inference using Typical RNA-Seq dataset (transcripts assembled using multiple assemblers) | 12 | 12.8 |
| DirInf-Pool |  | Direct Inference using Pooled RNA-Seq dataset (transcripts assembled using multiple assemblers) | 77 | 241.4 |
| DirInf-Orphan |  | Direct Inference using Orphan-rich RNA-Seq dataset (transcripts assembled using multiple assemblers) | 38 | 595.1 |
| BIND-Typical | Combined (BIND or MIND) | Direct-Inference using Typical RNA-Seq dataset plus BRAKER Typical predictions | 12 | 12.8 |
| MIND-Typical |  | Direct-inference using Typical RNA-Seq dataset plus Maker Typical predictions | 12 | 12.8 |
| BIND-Pool |  | Direct-Inference using Pooled RNA-Seq dataset plus BRAKER Pooled predictions | 77 | 241.4 |
| MIND-Pool |  | Direct-inference using Pooled RNA-Seq dataset plus Maker Pooled predictions | 77 | 241.4 |
| BIND-Orphan |  | Direct-Inference using Orphan-rich dataset plus BRAKER Orphan-rich predictions | 38 | 595.1 |
| MIND-Orphan |  | Direct-inference using Orphan-rich RNA-Seq dataset plus Maker Orphan-rich predictions | 38 | 595.1 |

**Figure 3.**
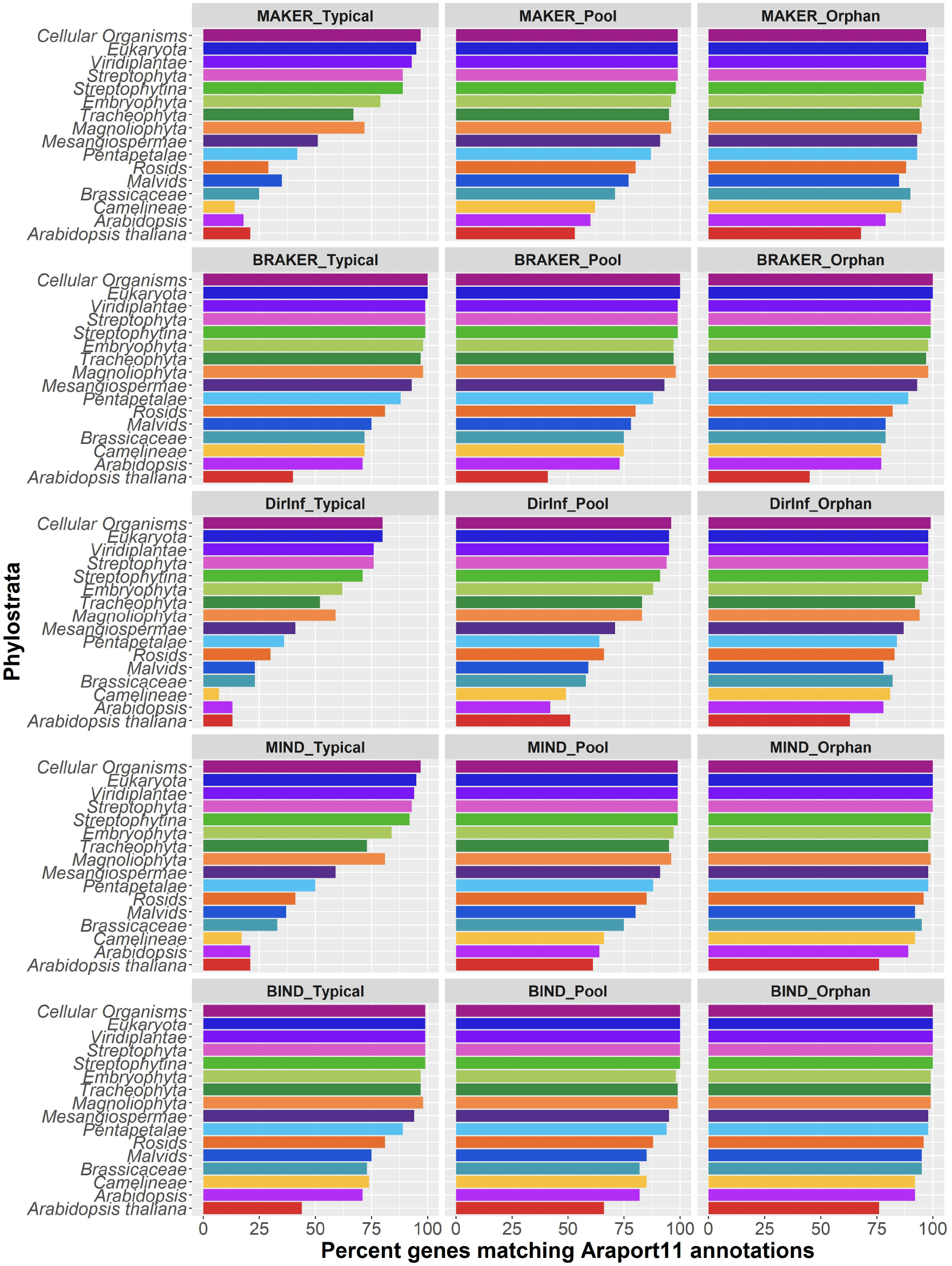
Arabidopsis Araport11-annotated gene predictions, shown by phylostratal designation. (See **Supplementary Table 3** for predictions of all tested scenario). For each prediction scenario, the ability to predict genes was greater for the genes of oldest phylostratum (Cellular Organisms) and gradually decreased for the younger phylostrata. More annotated genes were predicted when pipelines were supplied with a more diverse dataset, an exception being BRAKER pipeline predictions. Overall, BIND with orphan-rich dataset as input predicted the most genes matching Araport11 annotation.

Predictions by MAKER were highly dependent on the training dataset supplied. Total predictions varied between 80% and 96% of the annotated genes. For example, 22,065 of the annotated genes were predicted when the Typical RNA-Seq dataset plus its predicted proteins were used as input, whereas 25,649 of the annotated genes of Arabidopsis were predicted with training data from the Pooled dataset plus its predicted proteins (**Figure 3** and **Supplementary Table 4-A**).

This sensitivity to input evidence was much more pronounced for genes of the younger phylostrata. Twenty-one percent of the annotated orphan genes were predicted if MAKER was provided the Typical RNA-Seq dataset, versus 53% predicted from the Pooled dataset, and 68% for the Orphan-rich dataset (**Figure 3** and **Supplementary Table 4-A**). Even when provided a “gold-standard gene set” comprised of all annotated genes and their proteins (including all orphans) as training data, only 77% of the annotated orphans were predicted by MAKER (**Supplementary Table 4-A**).

The greater the diversity of the RNA-Seq evidence provided to MAKER, the more novel genes MAKER predicted that did not match any gene annotation, for example, the Typical dataset predicted 4,035 novel genes (1,183 of which were inferred by phylostratigraphy to be orphans), while the Orphan-rich dataset predicted 13,657 novel genes (7,194 of which were inferred by phylostratigraphy to be orphans) (**Supplementary Table 5-A**).

### Arabidopsis gene predictions by BRAKER

BRAKER differs from MAKER in that it uses RNA-Seq alignments to the genome for unsupervised training of GeneMark-ES/ET (**Figure 2**). Then, BRAKER selects a *subset* of the predicted protein coding genes to train AUGUSTUS and predict genes. BRAKER^31^ predicted 96%-97% of all annotated genes (**Figure 3** and **Supplementary Table 4-A**). Regardless of the training dataset provided (**Supplementary Table 3-A**), gene predictions differed in quantity and performance by less than 2% (**Supplementary Tables 6-A and 7-A**).

BRAKER’s ability to predict genes was greatest for the genes of the oldest phylostratum (“Cellular organisms”) and progressively decreased for younger phylostrata (**Figure 3** and **Supplementary Table 4-A**). For example, when provided the Pooled dataset as training data, BRAKER predicted 100% of ancient genes that traced back to Cellular Organisms, but only 41% of the orphan genes (**Figure 3**); 8,204 of the genes predicted by BRAKER were not annotated in TAIR (1,498 of these encode orphans) (**Supplementary Table 5-A**).

### Arabidopsis gene predictions by Direct Inference

Prediction by direct alignment of transcriptomic evidence to the genome is rarely used to annotate genes in newly sequenced genomes (e.g., **Supplementary Table 1**)^37^. That said, the use of cDNA and EST evidence-based prediction was a mainstay for early annotations^35,38^.

We considered that genes with non-canonical sequence features might be difficult to identify with machine learning algorithms. To mitigate this possibility, in addition to using transcript evidence indirectly as training data only, we developed an evidence-based approach that generates gene predictions, “Direct Inference” (Figure 2; Detailed in Methods). Briefly, the pipeline uses a genome-guided assembly, concatenates the transcripts, removes redundant transcripts, and determines ORF(s) in each inferred transcript. Because this approach directly relies exclusively on RNA-Seq alignments, only those RNAs that are expressed under the conditions sampled will be detected. Thus, we anticipated that providing RNA-Seq evidence collected from a wide variety of developmental stages, tissues and environmental conditions would be critical to maximize predictions when using Direct Inference. The overall performance of Direct Inference improved with larger dataset size **Supplementary Table 6-A**. Specifically, the diverse Orphan-rich dataset predicted nearly 96% of all annotated genes, and 63% of annotated orphans, while the Typical dataset predicted about 71% of all annotated genes, and 13% of the orphans (**Figure 3**, **Supplementary Table 4-A**).

### BIND and MIND: gene predictions combining *ab initio* (BRAKER or MAKER) with Direct Inference

Because *ab initio* methods can identify very high proportions of the conserved genes, while Direct Inference can identify genes that are represented in the RNA-Seq data evidence without regard to canonical structure or homology to genes in other organisms, we evaluated whether combining both approaches would maximize gene predictions across phylostrata **(Figure 2)**.

BIND increases the number of genes matching annotated genes compared to BRAKER or Direct Inference alone; MIND performed better than MAKER or Direct Inference alone (**Figure 3** and **Supplementary Table 4-A**). Using the Orphan-rich dataset as input, BIND, and to a lesser extent, MIND, predicted more annotated orphan genes than did either *ab initio* predictor alone. Base level F1 scores for overall prediction performance were comparable for BIND and MIND (~75%). Regardless of the RNA-Seq data input, BIND predicted the most accurate representation of all genes, and young genes in particular (**Supplementary Table 6-A**). MIND predicted 18,114 genes that didn’t match any annotated genes; BIND predicted 14,739 such genes.

Multiple low-expression transcripts are predicted by the *ab initio* methods, MAKER or BRAKER (**Supplementary Figure 1**). To remove low expressed novel predictions from predictions using the Orphan-rich dataset, we combined all the annotated and novel transcripts predicted; then, we filtered those novel transcripts with low-expression (see methods). When using the orphan-rich dataset as input, ~ 89% of the transcripts annotated in Araport11 are predicted by each pipeline; 6% more of the Araport11-annotated transcripts are predicted by one or more pipelines (**Figure 4**).

**Figure 4.**
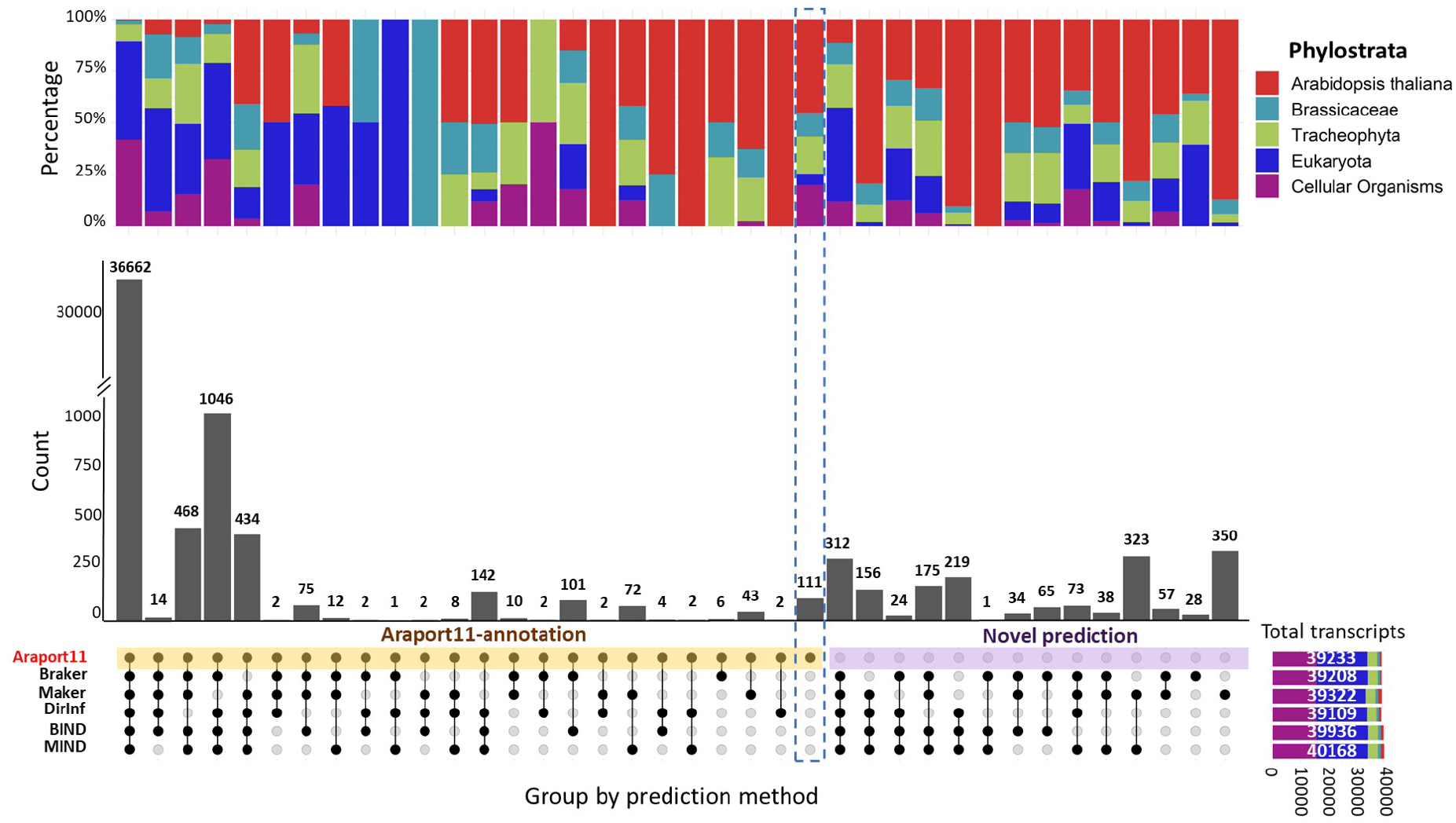
Upset plot of Arabidopsis protein-coding Araport11-annotated gene predictions and novel gene predictions for each pipeline. The Orphan-rich dataset was used as input data. The resultant 41,078 non-redundant predictions were filtered for low expression across over 5000 diverse RNA-Seq samples before plotting (see Methods and **Supplementary Figure 5**). Top panel, percentage of genes, binned by five phylostrata. Middle panel, numbers of predictions; Bottom panel, non-redundant genes grouped by prediction method; Bottom right panel, total number of genes in Araport11 and predicted by each pipeline, colored by phylostrata. When the Orphan-rich dataset is used as evidence, about 89% of all annotated genes are predicted by any single method alone.

Over 85% of annotated orphan genes were predicted by combining BIND and MIND, using the Orphan-rich dataset or the Pooled dataset. In contrast, using the Typical dataset, only 51% of annotated orphan genes were predicted by combining BIND and MIND, (**Supplementary Figure 2**).

We used ribosome footprinting data to assess the translation evidence for those genes predicted by BIND with the orphan-rich dataset. (See Supplementary Table 8A for all predictions) Predictions were filtered to remove those of low expression (see Methods). A limitation of this analysis was that the 185 Ribo-Seq samples that were publicly available did not represent diverse developmental and environmental conditions (**Supplementary Table 8B**).

Overall, 97% of the genes predicted by BIND had translation evidence. Translation evidence was greatest for proteins of the most ancient phylostratum (Cellular Organisms), decreasing for younger phylostrata (**Figure 5**). Ninety-eight percent of the predictions that matched annotated genes had translation evidence, while about 56% of the novel genes had translation evidence (**Supplementary Table 8-A**).

**Figure 5.**
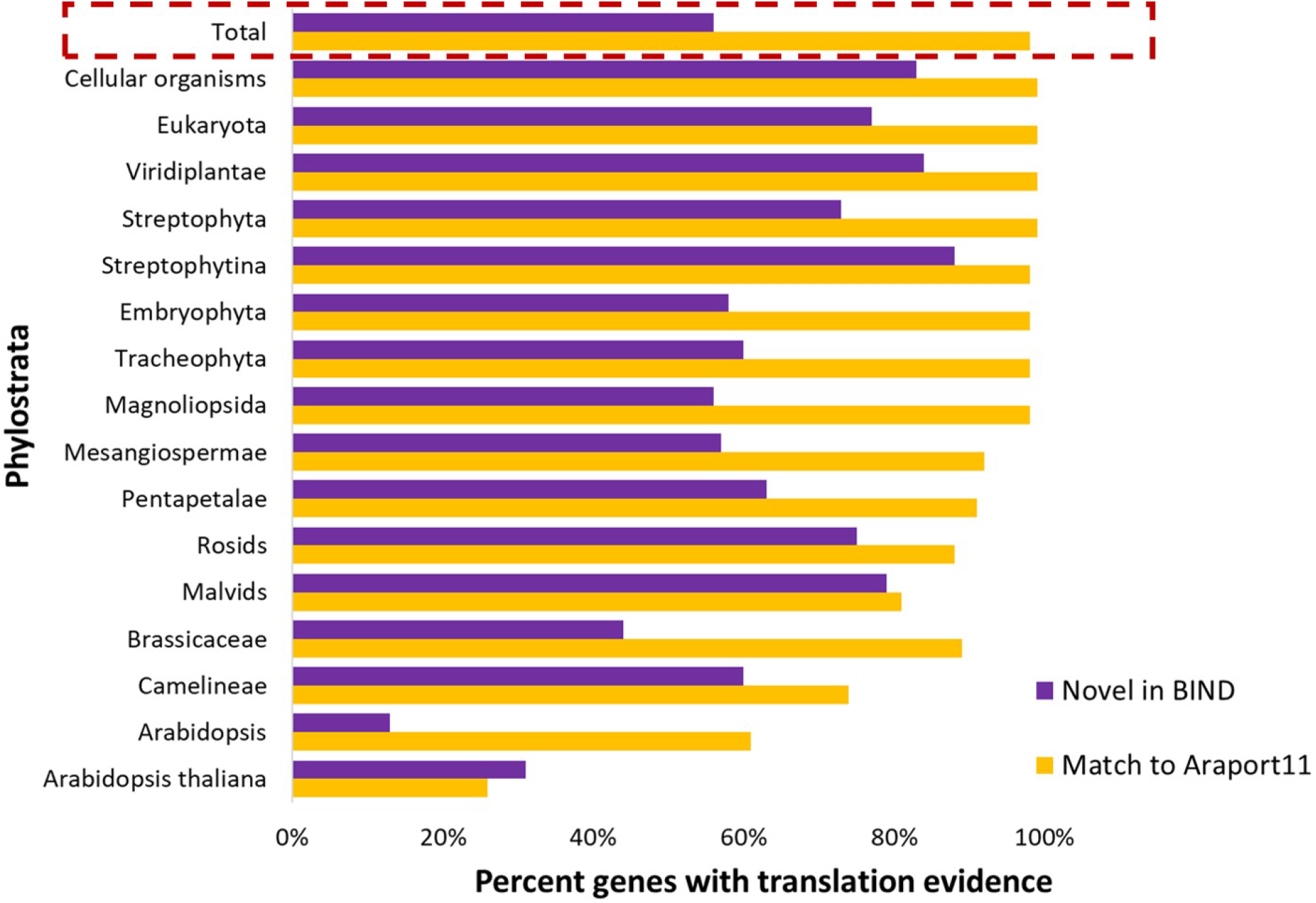
Translation evidence for BIND-predicted genes. Novel genes predicted by BIND were filtered for expression. Predicted proteins are binned by phylostratal designation. Translation signal was evaluated from the available ribosome profiling data. Translation evidence is consistent, but not sufficient, to indicate a protein-coding gene. Regardless of whether predictions matched to Araport11 annotations or were novel, younger genes had less translation evidence than ancient genes, as might be expected based on the sparser transcription patterns of younger genes.

### Application of the gene predictions scenario to other species

The identical approach was used to predict genes in the genomes of yeast and rice (**Supplementary Results**, **Supplementary Tables 4-7,9-11**, **Supplementary Figures 3,4**). For “Typical” size input data, the BIND approach provided improvement over any single method alone.

To enable BIND and MIND in a best practices format, we have implemented all Direct Inference core RNA-Seq processing steps in python using pyrpipe^34^ and the pipeline in Snakemake^39^ (see methods for details), we provide MAKER and BRAKER in singularity containers, and we have developed full, documented MIND and BIND pipelines in a versatile reproducible open source framework.

## Discussion

Structural annotation of genes in a genome is critical to making genomics data useful to the research community. However, the multiple protocols used to annotate genes, some of which exclusively rely on *ab initio* predictions; the wide variation in amount of training data input; and the dearth of reproducible methods (**Supplementary Table 1**) do not make for best-practice.

Quality gene annotation is particularly important for expression studies; this is because standard practice is to align RNA-Seq reads directly to reference transcriptomes consisting exclusively of annotated genes, or to align transcripts to the reference genome but count only the annotated genes. Both practices completely miss the detection of any genes that have not been annotated.

### Prediction of young genes is improved by a pipeline that combines ab initio/evidence-based methods and by supplying diverse RNA-Seq training data

By deploying the manually-curated, community-based gene annotations from the model species, Arabidopsis and yeast, as gold standards^35,38^ as well as the more recently annotated genes of Kitaake rice, we have explicitly illustrated some of the challenges of annotating young genes. The efficacy of every gene prediction scenario we tested, regardless of the pipeline used for annotation, the amount or quality of evidence supplied, or the species annotated, was strongly dependent on the phylostratal origin of the gene. As reflected by their lower visibility to *ab initio* annotation, younger genes appear to generally have a less recognizable sequence signature than do their more ancient counterparts, which have undergone hundreds of millions of years of selection. The clear trend is that the younger a gene is, the less likely it is to be predicted. This finding supports previous speculation^40^.

*Ab initio* pipelines infer genes by their structural motifs. In contrast, the Direct Inference pipeline uses the RNA-Seq evidence provided to directly infer genes. We show that combining *ab initio* predictions with evidence-based, direct inference of genes improves predictions compared to either method alone.

Our analyses reveal the extent to which selection of training data can significantly affect gene predictions by the MAKER, Direct Inference, MIND, and BIND pipelines. For these pipelines, extrinsic data from samples drawn from diverse environmental and developmental conditions improve the prediction of annotated genes. Although BRAKER is less affected by the quantity of training data, it has a ceiling on predictions. A possible explanation for this phenomenon is that, unlike MAKER, the extrinsic RNA-Seq evidence is filtered by BRAKER prior to making its final predictions. This filtering of the RNA-Seq evidence enables BRAKER to make reasonable predictions from even small RNA-Seq datasets, but also means it cannot take full advantage of the extra information offered by large, diverse RNA-Seq training datasets. The integration of Direct Inference predictions via BIND also removed most mis-spliced (artifactual) predictions made by BRAKER when annotating the transposon-rich rice genome (**Supplementary Table 9**). In the absence of extended diverse RNA-Seq data, BIND provides the best option for gene prediction.

If highly diverse RNA-Seq data is available, the BIND and MIND pipelines leverage this information. Including samples that often express high levels of young genes, such as reproductive tissues and stressed tissues^9,16,41–45^, is particularly critical for prediction of young genes.

### Challenges and limitations of gene prediction

Compelling evidence from bacteria, to yeast, to humans, indicates that many sequences that are not annotated as genes are transcribed and translated^3,26,27,46–52^. This “pervasive transcription” does not appear to occur randomly across genomes. Alignment to the genomes of all transcripts (with or without ORFs) from the RNA-Seq data summed from the diverse developmental, environmental conditions and genetic perturbations used in this study, indicates about 17% of the yeast genome and 24% of the Arabidopsis genome is not detectably transcribed. (Thirty-five percent of the rice genome was not detectably transcribed, but this result is based on more limited RNA-Seq evidence).

A major quandary in any gene annotation pipeline, and especially for evidence-based pipelines like Direct Inference^48,53–55^, is to determine which of the many gene predictions to retain and which to filter out. Each scenario that we tested predicted thousands of genes that were not annotated. One approach to provide more evidence for each prediction of a protein coding gene is to obtain translation evidence (Ribo-Seq and proteomics). This approach is limited by the available translation data, which may be non-existant for newly-sequenced species. In this study, over half of the *unannotated* ancient genes predicted by BIND for Arabidopsis also have translational evidence–even though the samples represented in the available Ribo-Seq data were limited as to the diversity of conditions represented.

Indeed, the same massive high-throughput sequencing that has enabled the identification and functional characterizations of genes has raised havoc with the traditional definition of a “gene”^3,5,48,55^. Perhaps most problematic for annotating genes is that genes themselves are on a continuum of “gene-ness” (**Figure 1**). A given gene may emerge, remodel, and recede in a continuum across evolutionary time.

Thus, some novel gene predictions might be “transcriptional noise”^26^ - fodder for *de novo* evolution of protein-coding genes; others might be ncRNAs with a non-translated or non-functional ORF^3,5,27,56^. Other novel gene predictions may be on the continuum between orphan gene and non-genic transcript or the path from gene to pseudogene^3,5^. And yet other novel gene predictions might be bonafide functional genes. No clear criteria articulate these products of the dark transcriptome. However, a first step is to annotate them, and provide evidence-based metadata.

Most protein-coding orphans appear to have arisen *de novo* from non-protein coding sequence, although various scenarios of defunctionalization and refunctionalization of existing genes provide another origin^1,3,4,27,57^. These youngest, most recently-formed protein-coding genes, encoding proteins with no amino acid sequence similarity to proteins of any other species, are among the least likely to have been functionally characterized^2^. A few orphan genes will persist over vast evolutionary time- as newer, younger genes arise in the organism the former orphans will become more and more ancient. Such genes will trace to deeper evolutionary origins (e.g., in the extant species Arabidopsis a gene that arose as an orphan at the inception of it’s Magnoliophyta ancestor would be classified in the Magnoliophyta phylastratum). About two thirds of the annotated genes of Arabidopsis trace back to a very early eukaryotic or a prokaryotic origin^2^. The most complex molecular process, catalysis appears predominantly restricted to genes of these ancient phylostrata^2^.

These considerations have resulted in a surprising diversity of definitions of a “gene” in current literature; an entire philosophy surrounds the concept of the nature of a gene (see **Supplementary Table 13** and references therein). To be consistent with current understanding of coding and non-coding genes, we will consider a gene as “*an expressed, heritable, DNA sequence that confers a chemical, developmental, morphological or biochemical phenotype under one or more conditions*”. This definition requires function, an expressed product (RNA or protein), and heritability. It also emphasizes the context-dependency of some genes–a sequence might confer a function that is essential only under particular conditions (e.g., exposure to a draught or a novel virus), or contribute a minor survival advantage^53^. An expressed sequence would be considered a protein-coding gene if it is translated and its protein product effects a phenotype under some condition(s), and hence under that condition it would be under selection (heritable). Under conditions in which function and heritability are weak, sequence evidence of selective constraints could be muddied. Because many sequences are in transition to (or from) “gene-ness”, this murkiness is inescapable.

### Application of gene annotation

In practice (see **Supplementary Table 1**), gene annotation is often biased against the annotation of young protein-coding genes. Annotation protocols that contribute to bias against young genes include: 1) Using RNA-Seq evidence from only a few conditions, which may not detect sparsely-expressed genes; 2) Relying exclusively on *ab initio* predictions, which miss many annotated orphans and other young genes; 3) Filtering out gene predictions that have only one exon, thereby excluding the many orphan genes with a single exon; 4) Removing gene predictions that do not have homologs in other species, which eliminates all orphan genes; 5) Removing predicted genes based on low overall expression or have many missing values, which eliminates the many orphan genes are expressed only under limited conditions. Thus, although these filtering strategies may minimize false positives, they also can exclude thousands of bonafide genes.

The challenge of confirming that young genes are annotated efficiently is not addressed by evaluating the extent that the annotation identifies Benchmarking Universal Single-Copy Orthologs (BUSCO, http://busco.ezlab.org), highly-conserved genes among the gene annotations. This method, though highly useful in determining how well a pipeline annotates more ancient genes, does not capture the efficacy of a pipeline in identifying orphans or other young genes. Recently, Scalizati et al^58^ have developed a benchmarking approach that takes phylostrata into account; herein we provide a different approach to benchmarking that considers phylostrata, and enables customization of phylostrata to include line-specific genes.

There is no substitute for manual curation in providing gene annotations that are useful to the community. An inclusive approach to annotating genomes that would benefit both curators and researchers would be to make accessible in the annotations the confirmed, curated genes along with predicted genes inferred based on expression and/or *ab initio* analysis, together with the available evidence for each. Thus, the signal of each predicted gene would be retained in the reference annotations. To minimize gene predictions that are actually “transcriptional noise”, filtering predictions based on transcript accumulation may provide a useful approach; however, applying a cutoff should take into consideration the RNA-Seq data that is available as well as the sparsity of expression of many orphan genes. Ultimately, providing broad, straightforward access to metadata on predicted genes will facilitate understanding of genome evolution and gene function.

The importance of retaining a broad view of gene expression is highlighted by the crucial functions that have been experimentally demonstrated for proteins encoded by both annotated and unannotated orphan genes. This is true particularly in the realm of orphan genes that confer resistance to abiotic and biotic stresses. Such genes may provide disruptive genetic elements that fundamentally reposition metabolic and regulatory networks^8,56,59,60^. The potential for transcripts that encode orphan proteins to be beneficial or essential has been reinforced by findings from synthetic biology research. This growing body of research reveals that even randomly-generated or evolutionarily-selected peptides with no clear similarity to any known proteins are often able to bind small molecules, such as ATP and amino acids, *in vivo*^61^. Furthermore, such “synthetic” orphan genes can have beneficial consequences, including developmental, stress-resistance, and longevity phenotypes, when expressed *in vivo*^61,62^. Thus, although a gene has only “recently” been subjected to selection pressure, it may be important to the organism. If, as we advocate here, information on predicted non-canonical genes was easily accessible, experiments could be designed to prioritize these inferred genes for experimental study and to elucidate the potential roles of these transcripts^34,39,63^ and to validate new pipelines by benchmarking against well-sequenced, well-annotated genomes. Furthermore, gene expression studies would include these predicted genes, and experimental biologists would gain a sense of how the genes might be acting.

### FAIR-ness of gene annotation protocols

Best practice for gene annotation is to use pipelines that can be easily reproduced and have been validated in model species. Unfortunately, neither have been standard practice (eg., **Supplementary Table 1**).

Two key factors to advance the field of gene annotation are *reproducibility* and the ability to modify the pipeline and its parameters. Our aim is implementation of Best Practice, reproducible pipelines for the methods we have deployed and developed in this research. To enhance reproducibility of the MIND and BIND pipelines, the workflow utilizes a package manager (for Direct Inference) and singularity containers (for MAKER and BRAKER) to install and run the bioinformatics tools. By automating the Direct Inference pipeline using pyrpipe^34^ and Snakemake^39^, we provide an end-to-end annotation solution that requires minimal user intervention.This automation, combined with extensive step-by-step documentation, enable a researcher aiming to annotate a novel genome to apply the methods to her/his own dataset.

We have facilitated future research in gene annotation by making all pipelines, containers, scripts, benchmark data, output data, and extensive step-by-step documentation open source and available (https://github.com/eswlab/orphan-prediction). Researchers can easily add/swap in new software for Direct Inference, and to some extent for BRAKER, and can optimize parameters of the software modules for Direct Inference and MAKER. A new pipeline can be compared to existing pipelines using benchmark data for model organisms, such as the TAIR and SGD gene annotations and the RNA-Seq data we provide. Researchers can compare and contrast the annotation pipelines by using the post-annotation analysis tools provided, or other tools selected by the researcher.

### Conclusion

We demonstrate that orphans and other young genes often elude annotation, and illustrate challenges and best practices in gene prediction. Our analyses showcase the importance of including diverse transcriptomic evidence and incorporating an evidence-based approach. BIND and MIND provide improved, user-friendly gene annotation, predicting genes for further study and curation. The BIND and MIND platforms will facilitate future research on gene annotation.

## Online Methods

### Software and data

Complete methods, all scripts used in this study, and all result files are documented, and a fully automated version of the direct inference pipeline implemented with pyrpipe and snakemake^39^ are available at https://github.com/eswlab/orphan-prediction. `pyrpipe` source code is available at https://github.com/urmi-21/pyrpipe. The `pyrpipe` package can be installed from bioconda (https://anaconda.org/bioconda/pyrpipe) or PyPi (https://pypi.org/project/pyrpipe).

### RNA-Seq, genome, and protein input data

*Arabidopsis thaliana Col0* genome (version Araport11) and reference genomes, GFF3 files, annotated transcript and protein sequences for (Araport11 version) were downloaded from The Arabidopsis Informatics Resource (TAIR)^35^. *Saccharomyces cerevisiae* analysis used genome (version R64-1-1) and annotated genes (version R64-1-1). *Oryza sativa* analysis used genome (GCA_009797565.1) and annotated genes (GCA_009797565.1) downloaded from NCBI.

We identified datasets of various sizes and complexities for Arabidopsis, yeast and rice (**See Supplementary Table 2, and Table 1**). The smallest datasets we term “Typical”; they are of sizes at the upper end of those often used in many gene annotation projects (The “Pool” datasets are about 10-fold larger than the “Typical” datasets, and are more diverse. The “Orphan-rich” datasets for Arabidosis and yeast are designed to maximize orphan representation. In developing this dataset, we reasoned that selecting samples which contain a breadth of orphan gene transcripts would be important because many, though by no means all, orphans are highly expressed under only a very limited set of conditions, such as a particular stress or a unique developmental stage^2,3,16^. The “Orphan-rich” datasets for *A.thaliana* and yeast were comprised of 38 RNA-Seq samples that each contained more than 60% of all annotated orphan transcripts. We compiled and predicted genes using three additional intermediate-sized datasets for Arabidopsis, and tested a “ground truth” dataset, composed of models of all Arabidopsis genes/proteins as annotated in Araport11 (“Ara11” dataset) (TAIR^35^) (**Supplementary Table 2-A**). RNA-Seq data and respective sample metadata were downloaded from NCBI-SRA as raw reads using the SRA-toolkit (v2.8.0)^64^ or automatically via pyrpipe^34^.

Protein sequence used as evidence in MAKER^30^ were generated in one of two ways: 1) For Arabidopsis, yeast, and rice, RNA-Seq reads were assembled using Trinity (v2.6.6)^65^, followed by open reading frame (ORF) prediction and translation using orfipy^29^ or TransDecoder (v3.0.1)^65^. 2) For Arabidopsis only, data was downloaded from Phytozome^66^ as predicted protein sequences for nine species: *Arabidopsis thaliana*, (*Glycine max, Populus trichocarpa, Arabidopsis lyrata, Conradina grandiflora, Setaria italica, Oryza sativa, Physcomitrella patens, Chlamydomonas reinhardtii*, and *Brassica rapa*).

### *Ab initio* annotation of genes by BRAKER

RNA-Seq raw reads were mapped to the indexed reference genome using HiSat2 aligner (v2.1.0)^67^ (default settings). The resultant SAM files were sorted and converted to BAM format. BAM files from each set of RNA-Seq samples were combined using SAMTools (v1.9)^68^ and provided as training for the BRAKER (v2.1.2) pipeline^31^, along with the unmasked genome. BRAKER is an automated pipeline to predict genes using GeneMark-ET (v4.33)^69^ and AUGUSTUS (v3.3.1)^70^. Briefly, GeneMark-ET is used to iteratively train AUGUSTUS, by generating initial gene predictions. GeneMark-ET-predicted genes are filtered and provided for AUGUSTUS training, followed by AUGUSTUS prediction, integrating the RNA-Seq information, to generate the final gene predictions (Figure 2, upper left panel).

BRAKER (v2.1.2) permits use of protein sequence training data to supplement the RNA-Seq training data. However, results with RNA-Seq plus protein evidence and with RNA-Seq evidence alone were virtually identical (the BRAKER User Guide also notes that it is not always best to use all evidence, https://github.com/Gaius-Augustus/BRAKER#running-braker). Thereafter, we used RNA-Seq evidence but not protein sequence for input training data to BRAKER.

### *Ab initio* annotation of genes by MAKER

To implement the MAKER (v2.31.10)^30^ pipeline, RNA-Seq data was assembled into a transcriptome using Trinity (v2.6.6)^65^; this CDS evidence, was supplied along with the unmasked genomes (Araport11). Depending on the case study and species, either CDS-only; CDS and translated proteins; or CDS and Phytozome proteins were supplied (**Supplementary Tables 2 and 3**).

MAKER was run in two successive rounds with default settings (**Figure 2**). In round one, transcriptome and protein data were aligned to the reference genome to generate crude gene predictions. These crude predictions were then used for training SNAP (release 2006-07-28)^71^ and AUGUSTUS (v3.2.1) *ab initio* gene predictors with default options. In round two, the Hidden Markov Models (HMM) for *ab initio* gene predictors, along with self-trained HMM of GeneMark-ES (v4.32) were used within MAKER to predict genes. MAKER finalizes the comprehensive sets of genes from all three predictors by ranking using Annotation Edit Distance (AED)^72^; the highest-ranking genes were retained for the final set of predictions. MAKER’s default output includes key metadata about gene predictions (evidence scores supporting each prediction, name of the component(s) within MAKER that generated the prediction).

### Evidence-based annotation of genes by Direct Inference

Raw RNA-Seq reads were assembled using three genome-guided transcriptome assemblers: *viz*. Class2, StringTie and CuffLinks^73–75^.

The BAM file generated by mapping reads to the Araport11-annotated indexed genome using HiSat2 (v2.1.0)^67^ was provided as training for the assemblers. The resultant assembled transcripts were used to predict ORFs using Transdecoder^65^ or orfipy^29^. We selected those complete ORFs over 150 nt. (Other user requirements might include: transcript length, number of exons, exon length, intron length, expression value, or presence of UTRs.) These data files, along with splicing junctions identified from the alignments using Portcullis^76^, were provided as input to Mikado^77^. Mikado pick was run with 28 threads, “nosplit” mode, report all orfs, and other default parameters.

We have engineered our Direct Inference pipeline in an automated, reproducible, scalable, and flexible manner by implementing the steps from downloading data through transcriptome assembly in the python library, pyrpipe^34^. Direct Inference requirements are relatively simple and are detailed in Singh et al.^34^.

Package managers, like Conda, or containers like Singularity or Docker, control for the execution environment and tool versions. We have implemented Direct Inference such that it can be used as is, or easily customized by the user. All requirements for the Direct Inference pipeline can be easily installed via Bioconda^78^ and the provided Conda environment file. Conda was built to install and update python packages and their non-python dependencies; it can also package software in other languages. The Conda environment enables a Direct Inference user to add or substitute software facilely to evaluate efficacy for different use cases. We selected Conda because it not only controls for the execution environment and tool versions, but it also works across platforms, and makes available a wide variety of bioinformatic software packages via the Bioconda channel. The Bioconda channel is a community driven repository to provide up-to-date bioinformatics software.

We used the Snakemake workflow manager^39^ to integrate the Direct Inference pyrpipe pipeline with the meta-assembly steps. The Direct Inference pipeline is available from https://github.com/eswlab/orphan-prediction/tree/master/evidence_based_pipeline.

### Combined gene predictions by MIND and BIND

The total predictions from Direct Inference were integrated with BRAKER predictions (BIND) or with MAKER predictions (MIND), using Mikado^77^ to merge predictions. The process and parameters of Mikado were identical to those for Direct Inference, except that the input files were changed to BRAKER (or MAKER) predictions and Direct Inference predictions. The merged predictions were finalized in GFF3 format.

### Implementation, computer allocations and ease of use: BRAKER, MAKER, and Direct Inference

**Supplementary Figure 7** gives an overview of BRAKER, MAKER, and Direct Inference prerequisites and features. For this comparison, we ran the three pipelines using the Arabidopsis genome with the Typical and Pooled RNA-Seq datasets as input (Table 1). Without a container, installation is far easier for BRAKER and Direct Inference than for MAKER.

To run Direct Inference with pyrpipe and Snakemake requires a single step - the input of SRA accession IDs; BRAKER requires a user to execute three single command-line operations. In contrast, MAKER is a hands-on program, in which a user must manually perform most steps, including transcript assembly, translation, evidence collection, training *ab initio* gene prediction programs (snap, AUGUSTUS, GeneMark). Users must also manually track outputs of all these steps and use them to run multiple iterations of MAKER.

Being able to change software and software parameters in annotation pipelines is important because genomes of different organisms have different characteristics, and different quantities of RNA-Seq data will be available depending on the species. Because of its modular construction in python, the Direct Inference pipeline can be modified with respect to the software program parameters. Furthermore, the software programs themselves can be exchanged or added. The RNA-Seq processing components implemented via pyrpipe are simple to modify. Snakemake provides multiple options for executing and scaling the pipeline on different HPC systems. Running MAKER entails a higher manual overhead, but this design aspect allows for the parameters to be changed by the user^79^ However, MAKER is not designed to enable changes of software programs. Because BRAKER is hard coded, changing parameters is difficult. However, some software programs that meet BRAKER’s core script requirements can be swapped or added.

Implementing computational pipelines that are easily reproducible can be a challenging task^34,63^. Because bioinformatics pipelines run a number of software programs that interact with the operating system libraries and with each other, controlling for the version, execution environment, and parameters of each program is essential for reproducible pipelines. We have implemented the Direct Inference pipeline keeping this principle in mind. All the required dependencies for the Direct Inference pipeline are automatically installed inside an isolated Conda environment. Centralized parameter management makes it easy to share and modify pipeline parameters. To maximize reproducibility for the MAKER and BRAKER pipelines, we have provided Singularity containers; these containers execute the tool in a virtual environment.

Gene annotation using large RNA-Seq datasets may best be done using multiple nodes. Because we implemented Direct Inference using the Snakemake workflow manager, it can be conveniently managed and scaled for multiple nodes. The BRAKER container is also optimized for use on multiple nodes. The MAKER container was not optimized for running on multiple nodes. For MAKER, the user would need to correctly configure the MPI program on both host and on the container-which would be quite challenging, and eliminate the benefits of having a container. Further, the configuration would likely require admin privileges on the HPC; general users rarely have such privileges.

If running the pipelines on a single node, relative efficiency is dependent on data size. When run with the large Pooled or Orphan datasets, BRAKER is more efficient than Direct Inference or MAKER in terms of disk usage, disk I/O (Input/Output). In contrast, when run with the “Typical” dataset (12.8 GB), Direct Inference is more efficient than BRAKER or MAKER in terms of disk usage.

### Comparing prediction scenarios and estimating performance of gene prediction methods

The results of each gene prediction pipeline scenario were compared to the existing annotations using Mikado Compare^77^. Gene structure annotation predictions were provided to Mikado as GFF3 files. Similarity statistics are reported for each gene locus individually. Performance calculations were based on sensitivity (Sn), precision (Pr), and the combined performance metric, F1 score^80^. Sn is a measure of the percent of predicted genes matched to all reference annotated genes (true positives) (Equation 1). Precision (Pr) is a measure of of the specificity of the predictions, that is percent of reference annotated genes (true positives) matching all predicted genes (Equation 2). The F1 score combines the sensitivity and precision as a measure of performance (Equation 3).

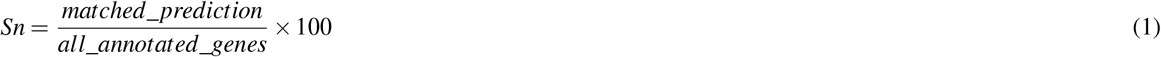

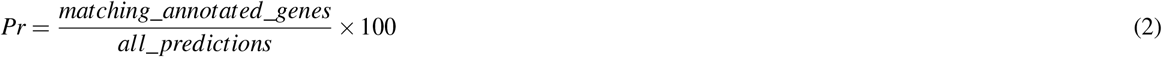

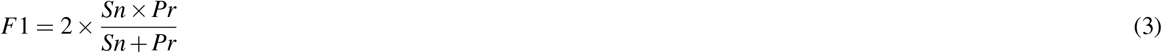

### Gene and transcriptome features for each prediction

The gene features were identified by Genome Annotation Generator (GAG)^81^ based on the final GFF3 files (available at https://github.com/eswlab/orphan-prediction/tree/master/prediction_gff3). These include: number and length summary for genes, mRNAs, exons and introns; genome coverage by CDSs and genes; overlapping and contained genes in **Supplementary Table 9**. The coverage of transcribed sequence was calculated by bedtools2^82^ based on the predicted transcriptome.

### RNA-Seq expression analysis

To investigate the expression level for *A.thaliana* predictions, we collected 5,210 RNA-Seq samples from NCBI-SRA database. Criteria were: paired end reads (layout) and transcriptomic (source) and rna-seq (strategy) and illumina (platform) and organism (Arabidopsis thaliana). Salmon^83^ (mapping-based mode) was used to quantify the expression of all genes predicted by orphan-rich dataset and annotated in Araport11. We calculated the median of mean expression of all Araport11 genes across all samples; this median was 2.18 tpm. Then, those unannotated transcripts with expression of at least 2.18 tpm in at least 100 samples were used to plot *Figure 4*. The less-expressed transcripts were considered as low-expressed transcripts. The expression for all predicted transcripts was visualized by R and shown in **Supplementary Figure 1**.

### Ribo-Seq analysis

To investigate the translational activity of BIND predictions, we analysed 185 samples of Ribo-Seq data from 21 studies (**Supplementary Table 8**). Raw reads were downloaded from NCBI-SRA, and the SRA-toolkit (v2.8.0) was used to convert the raw reads to a FASTQ format. BBDuk was used to remove adapter sequences from the 3énd of reads, and rRNA reads were identified and removed using BBMap^84^. The cleaned Ribo-Seq reads were aligned to the reference genome by STAR aligner (v2.5.3)^85^. Ribosome-bound ORFs were detected and quantified by Ribotricer^86^, which considers the periodicity of ORF profiles, using the recommended parameters for Arabidopsis. A gene with at least five codons with non-zero reads was considered to have translation signal in a Ribo-Seq sample^86^.

### Phylostratigraphy

Phylostratigraphic analysis based on the homology of predicted proteins to proteins in clades of increasing depth (age) was inferred using the R-platform *phylostratr* software (v0.2.0)^13^. The focal species were set as “3702” for *A. thaliana*, “4932” for *S. cerevisiae*, and “39947” for *O. sativa*. A database of protein sequences was created by *phylostratr* from hundreds of species. The databases of protein sequences for Arabidopsis, yeast and rice were created by *phylostratr* from diverse species, including custom species selected based on evolutionary clade and genome/proteome quality. The customized, more comprehensive, set of proteins was provided for several species, and protein sequences of the remaining species were downloaded as additional Uniprot (**Supplementary Table 14**). The phylostratal designations for Arabidopsis have been annotated in TAIR’s jbrowse “phylostratigraphy identification” track (arabidopsis.org). We inferred the age of proteins from the ORFs of the genes that were predicted by five annotation pipelines using different datasets for three test species (**See Supplementary Tables 7, 11 and 12**) and used these assignments to benchmark each gene prediction pipeline for its efficacy in predicting genes according to their inferred phylostrata.

## Supporting information

Supplementary Results

## Data availability

RNA-Seq used as training data input to *ab initio* predictors and for Direct Inference alignments are specified in **Supplementary Table 2 and 3** and were obtained from National Institutes of Health Sequence Read Archive https://www.ncbi.nlm.nih.gov/sra. Gene predictions for each scenario and species are provided as GFF files at https://github.com/eswlab/orphan-prediction/tree/master/prediction_gff3. The full phylostratal designations and other metadata for annotated and predicted genes in Arabidopsis and rice are provided in **Supplementary Tables 11-12**.

## Code availability

Overview with links to source code is available at https://orphan-prediction.readthedocs.io/en/latest/; recommended for analyses. All source code including for pipelines, downstream analyses, and creation of figures is available and documented at https://github.com/eswlab/orphan-prediction.

## Acknowledgements

We are grateful to Nathan Weeks, USDA Agricultural Research Service, for development of singularity containers for MAKER and BRAKER. We thank Ethalinda Cannon, USDA Agricultural Research Service, for her prescient suggestions on gene metadata, Andrew J. Severin, Basil J. Nikolau, and Gene Prediction Summit members for valuable discussion. Linor Vaknin kindly provided expert design consultation on Figure 1. This research was supported in part by the Center for Metabolic Biology and by the US. Department of Agriculture, Agricultural Research Service. USDA is an equal opportunity provider and Employer. Mention of trade names or commercial products in this publication is solely for the purpose of providing specific information and does not imply recommendation or endorsement by USDA. This work used Extreme Science and Engineering Discovery Environment (XSEDE)(National Science Foundation Grant No. ACI-1548562) via Bridges HPC environment allocation TG-MCB190098 and Iowa State University cyberinfrastructure. This study is based upon work supported by the National Science Foundation under Grant No. IOS 1546858 to ESW.

## Author contributions statement

JL designed and performed Arabidopsis, yeast and rice analyses, analyzed the data, developed code, and wrote the paper; AS designed Arabidopsis analyses, developed code, and wrote the paper. ESW conceived the study, contributed annotation and analysis, and wrote the paper; PB processed RNA-Seq data for Arabidopsis; US designed MIND and BIND with reproducibility assurances and revised the paper; JC beta-tested the annotation pipelines and provided documentation; ZA performed parts of the phylostratigraphic analysis. All co-authors wrote the parts of the paper they contributed to and commented on the entire paper.

