## Supplementary Figures 1-7 for "Foster thy young: Enhanced prediction of orphan genes in assembled genomes"

**Note:** Scripts for all supplementary figures can be found at https://github.com/eswlab/orphan-prediction/tree/master/plots_publication

Supplementary Figure 1. **Expression of Araport11-annotated Arabidopsis genes and genes predicted by each annotation pipeline.** The Orphan-rich dataset was used as input data to the pipelines. Each expression heatmaps shows expression from 5210 RNA-Seq samples. Y-axis, samples are sorted by expression, independently for each gene; highest expression is on left-hand side of each panel. X-axis, genes; genes are divided into three phylostrata (top panel, genes inferred as Cellular Organisms to Tracheophyta; middle panel, genes inferred as Magnoliophyta to *Arabidopsis*; bottom panel, genes inferred as orphans (*Arabidopsis thaliana*)). **A**, genes matching Araport11-annotated genes; **B**, novel predicted genes, not annotated in Araport11.

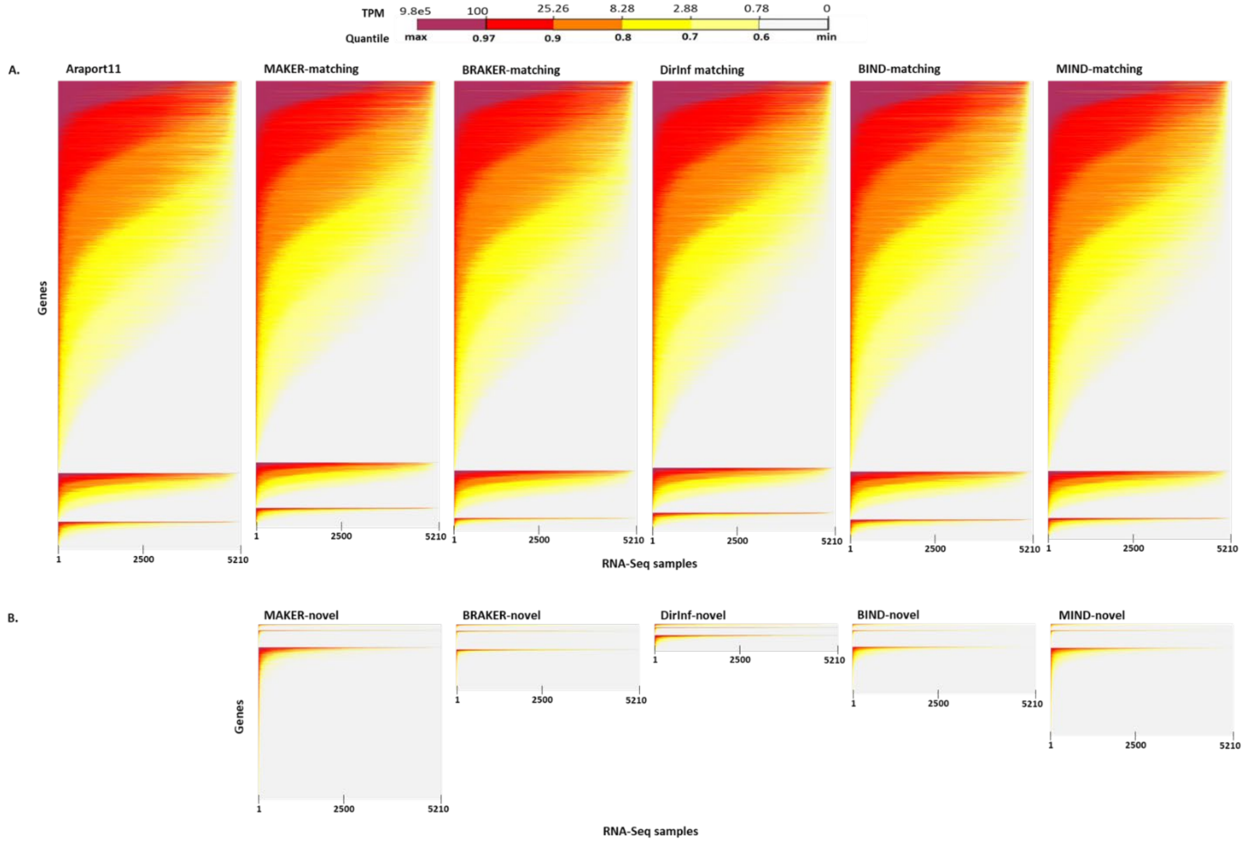

Supplementary Figure 2. **Arabidopsis orphan genes: Upset plots showing Araport11-annotated orphan genes and novel predicted orphan genes identified by each prediction scenario. A.** Orphan-rich dataset input. **B**. Pooled dataset input. **C.** Typical dataset input. **A-C.** top panel, number of genes in each group; bottom panel, non-redundant orphan genes group by prediction method; bottom far right bar chart, total number of orphan genes predicted by each method and annotated in Araport11. Dashed rectangles, annotated genes not predicted by any method.

**A. Araport-11-annotated and novel orphan transcripts (8,214) predicted using Orphan-rich dataset.**

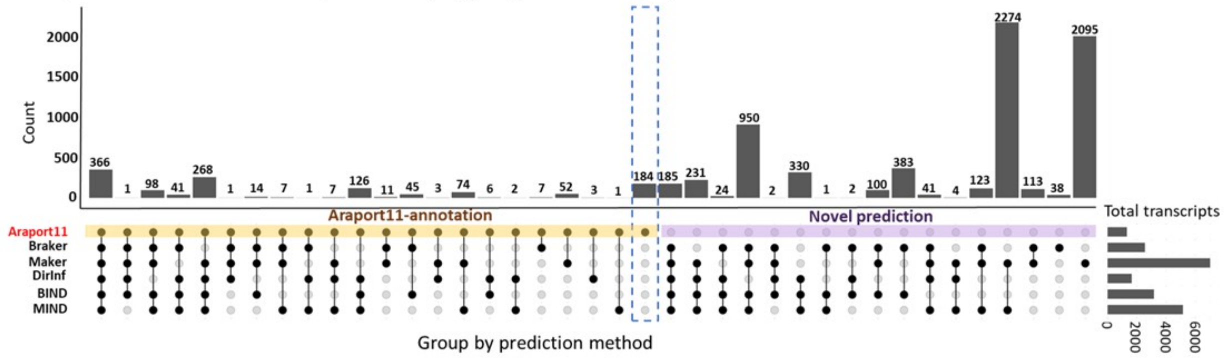

**B. Araport-11-annotated and novel orphan transcripts (5,660) predicted using Pooled dataset.**

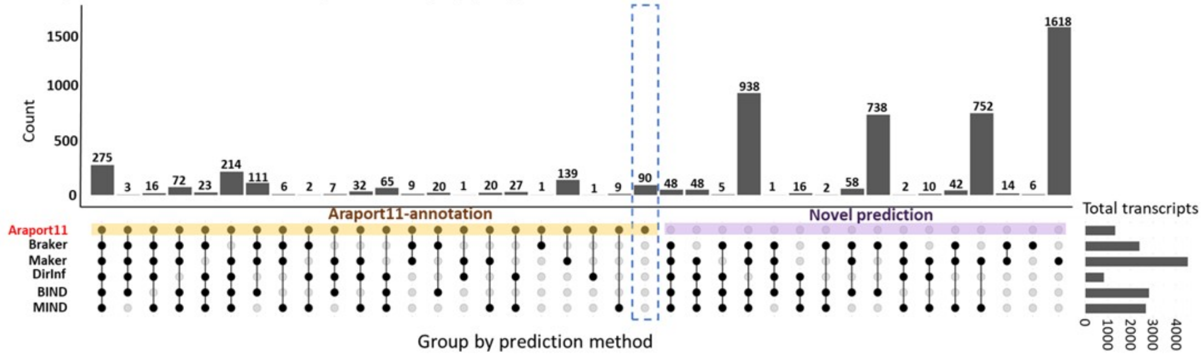

**C. Araport-11-annotated and novel orphan transcripts (1643) predicted using Typical dataset.**

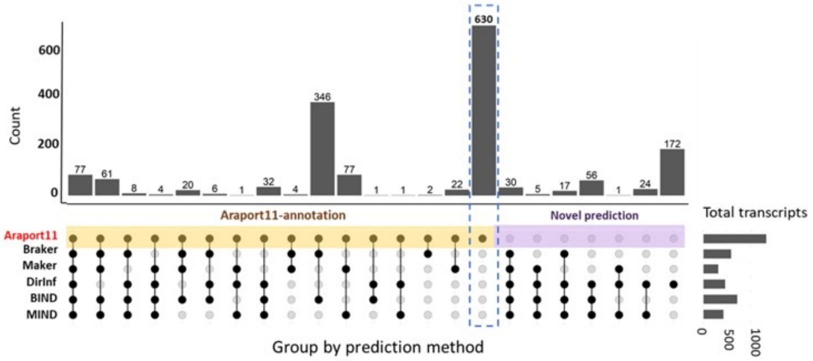

Supplementary Figure 3. **Yeast orphan genes: Upset plots showing SGD-annotated orphan genes and novel predicted orphan genes identified by each prediction scenario.** Unlike for Arabidopsis, most SGD-annotated yeast orphan genes were not predicted, even with the most diverse RNA-Seq data as input. The Orphan-rich dataset gave the highest numbers of SGD-annotated orphan gene predictions, about double that for the Typical dataset. **A.** Orphan-rich dataset input. **B**. Pooled dataset input. **C.** Typical dataset input. **A-C.** top panel, number of genes in each group; bottom panel, non-redundant orphan genes grouped by prediction method; bottom far right bar chart, total number of orphan genes predicted by each method and annotated in SGD. Dashed rectangles, annotated genes not predicted by any method.

**A. SGD-annotated and novel orphan transcripts (496) predicted using Orphan-rich dataset.**

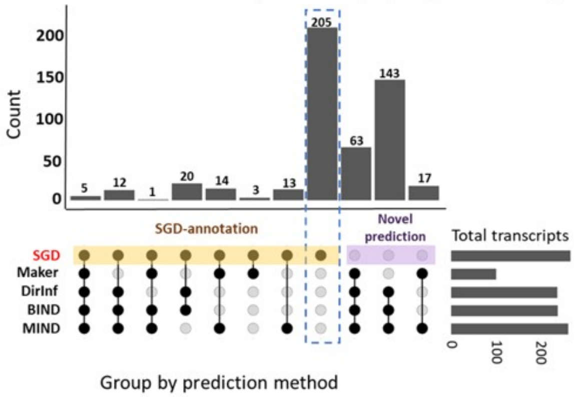

**B. SGD-annotated and novel orphan transcripts (425) predicted using Pooled dataset.**

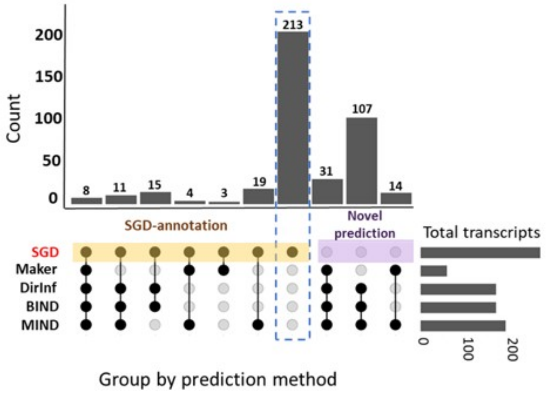

**C. SGD-annotated and novel orphan transcripts (286) predicted using Typical dataset.**

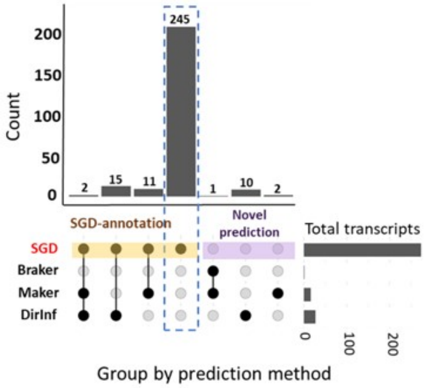

Supplementary Figure 4. **Rice orphan genes: Upset plots showing NCBI-annotated orphan genes and novel predicted orphan genes identified by each prediction scenario.** In rice, BRAKER predicts multiple mis-spliced orphan genes; BIND removes most of these. **A**. Pooled dataset input. **B.** Typical dataset input. **A-B.** Top panel, number of genes in each group; bottom panel, non-redundant orphan genes grouped by prediction method; bottom far right bar chart, total number of orphan genes predicted by each method and annotated in NCBI. Dashed rectangles, annotated genes not predicted by any method.

**A. NCBI-annotated and novel orphan transcripts (31,324) predicted using Pooled dataset.**

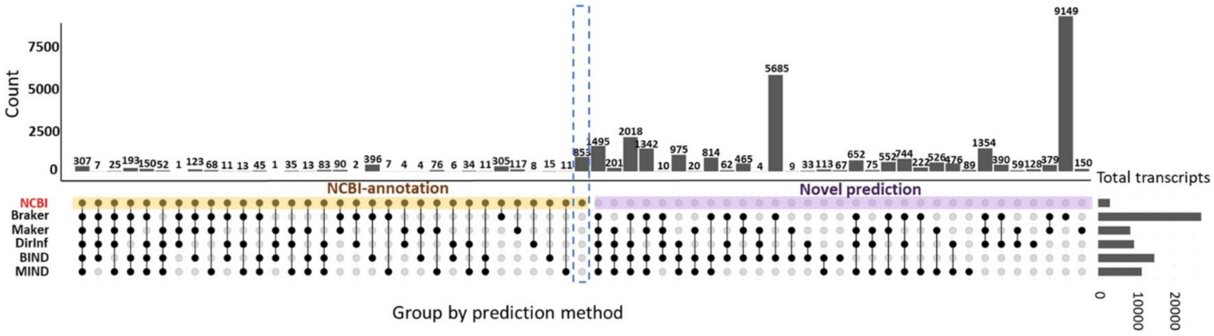

**B. NCBI-annotated and novel orphan transcripts (26,794) predicted using Typical dataset.**

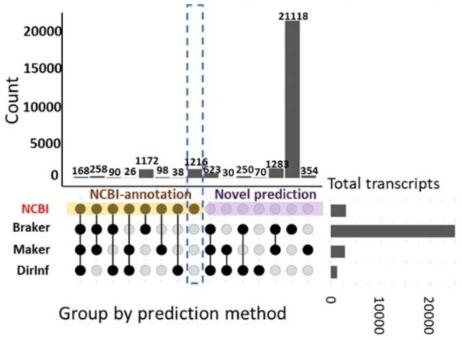

Supplementary Figure 5. **Upset plot of all Arabidopsis protein-coding Araport11-annotated genes and novel genes for each prediction pipeline.** The Orphan-rich dataset was used as input data. Top panel, percentage of genes if divided by five phylostrata. Middle panel, number of predictions; Bottom panel, non-redundant genes (63,081 in total) grouped by prediction method; Bottom right panel, total number of genes in Araport11 and predicted by each prediction method, colored by phylostrata.

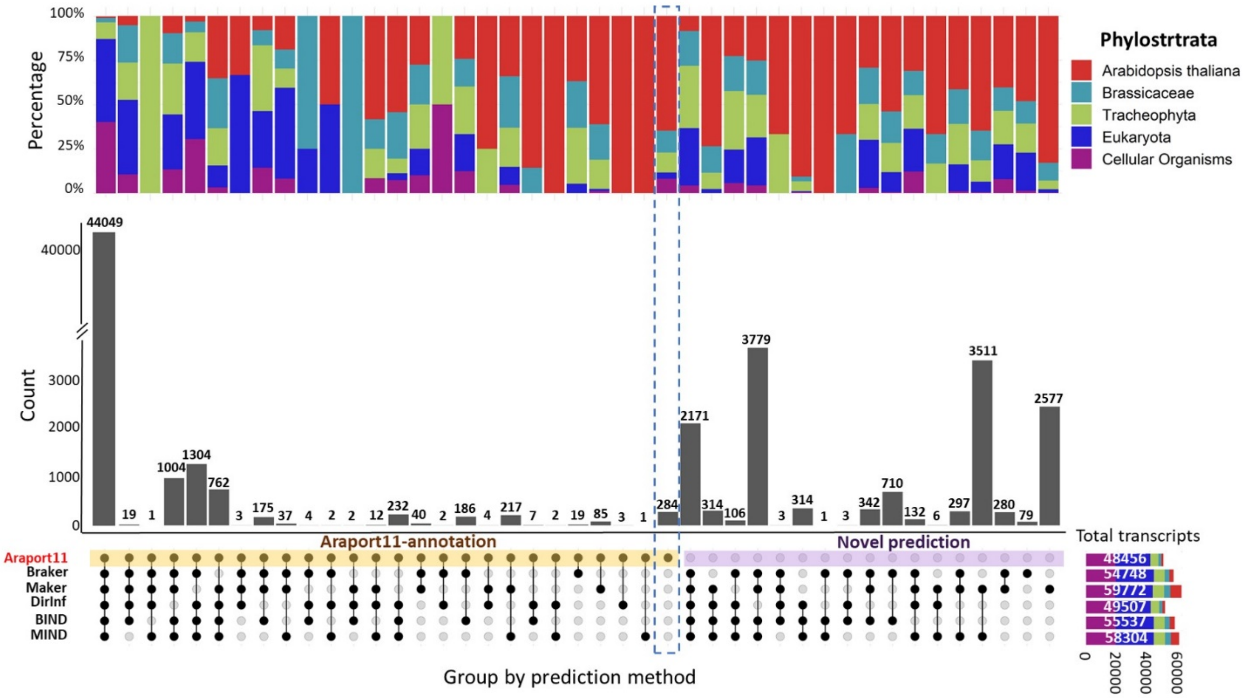

Supplementary Figure 6. **Translation evidence for all BIND-predicted genes (match to Araport11 vs novel in BIND) of Arabidopsis.** Translation signal was evaluated from the ribosome profiling data available for Arabidopsis; these samples were derived from somewhat limited conditions (See Figure 5 in main text for highly expressed prediction only, and Ribo-Seq samples in Supplementary Table 8-B). Predicted proteins are binned by phylostratal designation. Regardless of whether genes matched to Araport11 annotation, the younger genes had less translation evidence than ancient genes, as might be expected based on their sparse transcription patterns.

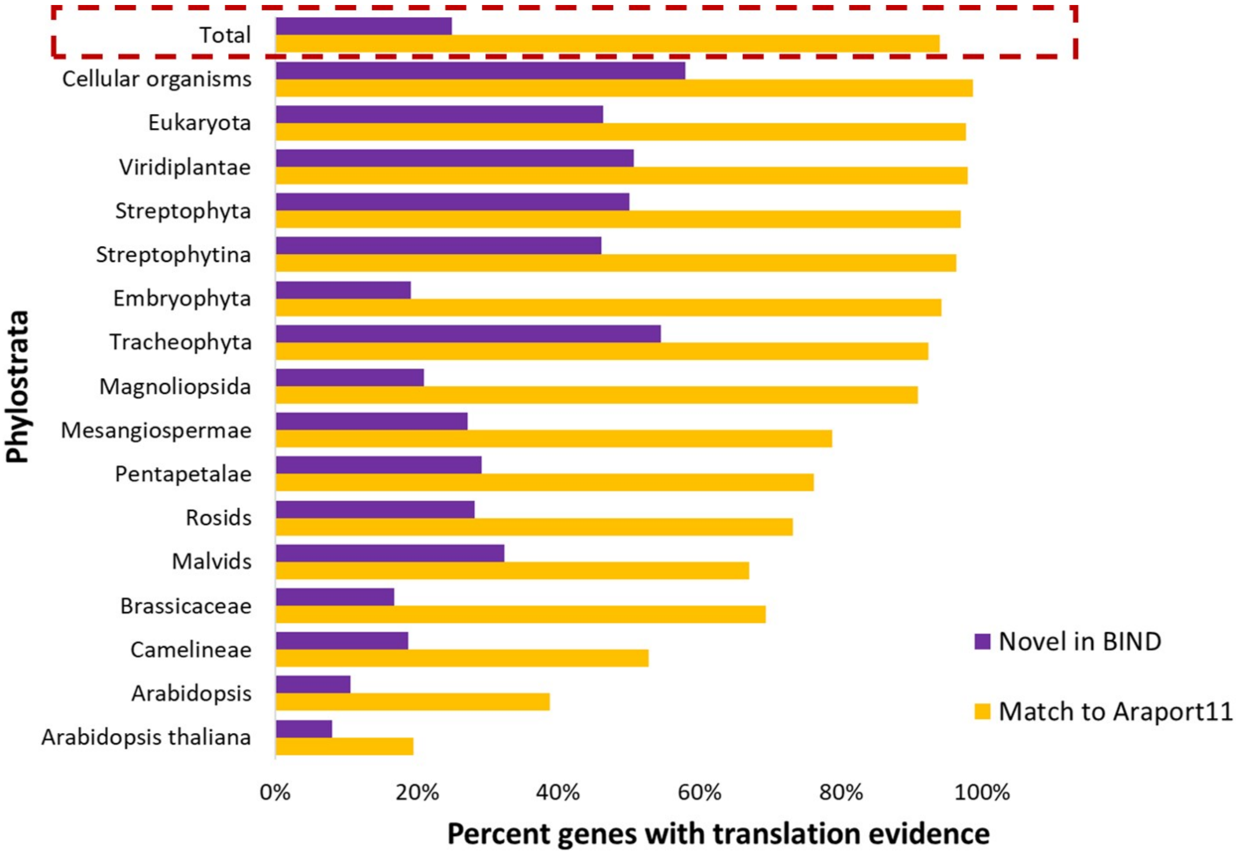

Supplementary Figure 7. **Overview of BRAKER, MAKER, and Direct Inference prerequisites and features.** For this comparison, we ran the three pipelines using the Arabidopsis genome with the Typical and Pooled RNA-Seq datasets as input (as detailed in Table 1 in manuscript).

**Installing the software** (Installation was tested by a bioinformatician with 10 years' experience, and independently by a genetics PhD student with two years of bioinformatics experience. Without containers, both researchers were able to install BRAKER and Direct Inference pipelines in less than 30 min. In contrast, MAKER has numerous dependencies that need to be installed independently and then configured, before compiling MAKER. Thus, MAKER installation took more than 20 hours for the highly experienced bioinformatician (not including testing with a toy dataset), and the less experienced researcher was not able to install MAKER even after multiple attempts. With containers, as supplied in the pipelines, the process of installation is obviated.

**Number of prerequisite software programs and perl modules:** for MAKER (either as dependency or as tool), 13 programs and 11 perl modules; for BRAKER, 7 programs and 11 perl modules; for Direct Inference, 7 programs.

**Running the software:** Direct Inference takes a single step to run: inputting SRA accession IDs. BRAKER requires a user to execute three single command-line operations: downloading the SRA data, mapping the raw reads to the genome, and running BRAKER. In contrast, to run MAKER, a user must manually download SRA data, assemble transcripts, translate them, collect evidence, train gene predictors, conduct snap training by a series or sequential commands and using files collected from specific locations of the output, and run multiple iterations of AUGUSTUS.

**Flexibility of the pipelines:** The Direct Inference pipeline can be modified with respect to the software programs used and their parameters. The RNA-Seq processing components implemented via pyrpipe are simple to modify. Snakemake provides flexible options for executing and scaling the pipeline on different HPC systems.
The design of MAKER enables users to change parameters of its software programs (Wang et al., 2018). MAKER has fixed software programs that it needs to use for various steps. You cannot switch to a different program to do the same job. It is possible to update to different versions of the same program if it is compatible with the MAKER version. If major changes are made to the versions used. recompilation of the MAKER executable may sometimes be required.
BRAKER is mostly hard-coded (training GeneMark, gathering hints, training and predicting using AUGUSTUS) and users do not have options to edit the settings. Thus, BRAKER is simple to execute but hard to optimize for particular datasets or organisms. BRAKER does provide fixed choices for a user to swap software for some functions (such as protein alignment); however, the core programs (such as GeneMark and AUGUSTUS) are not designed to be altered.

**Disk usage:** For the Typical RNA-Seq dataset (12.8 GB), Direct Inference is more efficient than BRAKER or MAKER in terms of in terms of disk usage, disk I/O (Input/Output), when run on a single node.
For the Pooled RNA-Seq dataset (241 GB), BRAKER is more efficient than Direct Inference or MAKER in terms of disk usage, when run on a single node. (MAKER is slow because it creates millions of intermediary files by partitioning chunks of genome and writing predictions for each predictor as separate files, and sequences. Direct Inference is slow because it uses different assemblers including Cufflinks (Trapnell et al., 2012), which is slow. It was important to include Cufflinks in Direct Inference because we wanted to discover an exhaustive set of transcripts using most of the popular tools.)

**Containers:** Because we have implemented the Direct Inference pipeline into Snakemake workflow management system, it is simple to run Direct Inference rapidly using multiple nodes. The containerized version of BRAKER also runs facilely on multiple nodes.  However, the MAKER container is not optimized for running on multiple nodes, due to technical considerations.
MIND (and BIND) require the prerequisites and programs needed for MAKER (and BRAKER) added to those for Direct Inference.

| Feature | BRAKER | Direct Inference | MAKER |
|---|---|---|---|
| Installation | Moderate installation requirements if not using the containers | Moderate installation requirements if not using containers | Heavy installation requirements. Available container is not optimized for MPI (to run on multiple nodes) |
| Installation with container or package manager | Yes, available via Singularity | Yes, all dependencies are available via Conda & Singularity | Yes, available via Singularity (without MPI) |
| # Interventions by human user to run pipeline | 3 | 1 | 11+ |
| Interventions by human user to run pipeline | 1. Provide SRA ACCESSION<br>2. Download data<br>3. Run BRAKER | 1. Provide SRA ACCESSION | 1. Provide SRA ACCESSION<br>2. Download data<br>3. Assemble (trinity or other assembly method) into ESTs<br>4. Use TransDecoder to generate proteins<br>5. Configure MAKER CTL files to use ESTs, proteins, and turn-off ab initio predictions<br>6. Run MAKER<br>7. Generate gff3, proteins, transcripts (first round)<br>8. Use genome file to train genemark<br>9. Filter gff3 and use to train SNAP (Requires *multiple* user interventions) Use filtered gff3 to train AUGUSTUS (Requires multiple user interventions)<br>10. Modify the CTL files to use the first round GFF3 (saves some time), and enable ab initio predictions (use profiles generated from training AUGUSTUS)<br>11. Generate gff3, proteins, transcripts (second round) |
| Ability to modify parameters | No, parameters for the programs running within BRAKER are inaccessible | Yes, parameters are defined in yaml files and are easy to modify | Yes, via CTL files |
| Able to substitute/swap in new software | Yes, swappable using variables as long as they meet BRAKER (main script) requirements | Yes, programs work in modules and are not interdepdendent | No, Updating programs requires re-building MAKER |

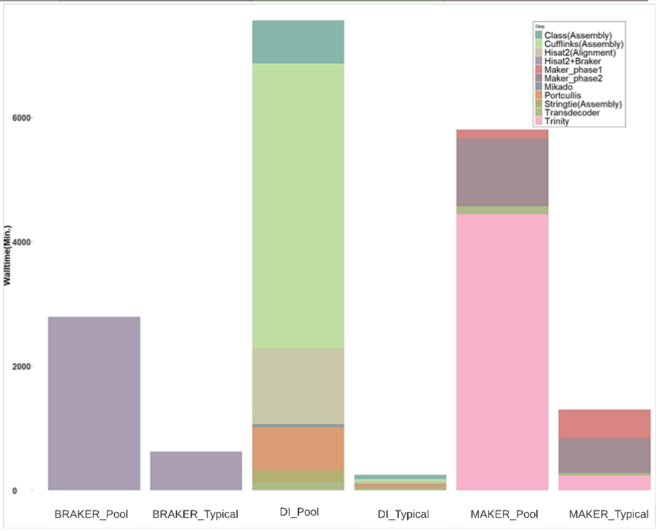
