## Supplementary Results for "Foster thy young: Enhanced prediction of orphan genes in assembled genomes"

### Gene predictions for *Saccharomyces cerevisiae*

We evaluated the efficacy of the gene prediction pipelines on a disparate genome, that of the fungus, *S. cerevisiae*, using the highly curated Saccharomyces Genome Database (SGD)[1] gene annotations for benchmarking. As for Arabidopsis, yeast genes have been manually annotated through experimental evidence over tens of years. We assembled three datasets of varied sizes and compositions: a "Typical" dataset; a "Pooled" dataset, consisting of the "Typical" dataset plus other RNA-Seq data from samples from varied conditions; and an "Orphan-rich" dataset, comprising 38 RNA-Seq samples (selected from 3,457 high-quality samples[2]) that are highly represented in SGD-annotated orphan genes (**Supplementary Table 2-B and 3-B**). We partitioned the SGD-annotated genes according to the phylostratigraphic inferences from[2].

Yeast genes were predicted using the MAKER, BRAKER, Direct Inference, MIND, and BIND gene annotation pipelines, each in combination with the "Typical", Pooled" and "Orphan-rich” datasets as extrinsic training data, thus providing a total of 15 gene annotation scenarios.

MAKER’s ability to predict genes was greater for more ancient genes (e.g., Cellular Organisms and Fungi, PS1-4) than genes of younger phylostrata (*Saccharomyces* PS10, and orphans PS11), regardless of the extrinsic evidence provided (**Supplementary Table 4-B**). MAKER predicted more annotated genes when the Pooled or Orphan-rich data was provided as input, matching 74% and 71% of SGD-annotated genes, respectively.

BRAKER’s ability to predict genes was greater for genes of more ancient phylostrata than those of younger phylostrata (**Supplementary Table 4-B**). BRAKER yielded predictions that differed in quantity and F1 score by less than one percent with each of the three datasets. BRAKER predicted 82% of all SGD-annotated genes (Supplementary Table 4-B) with an F1 score of 97% (Supplementary Table 6-B, base level). However, BRAKER did not predict a single orphan gene in yeast, even when supplied the Orphan-rich dataset.

Using the Orphan-rich dataset as input, the Direct-Inference pipeline predicted nearly 83% of all annotated genes and 13% of SGD-annotated orphans. In contrast, using the Typical dataset, Direct-Inference predicted only 33% of all SGD-annotated genes and 6% of the annotated orphan genes (**Supplementary Table 4-B**).

BIND and MIND, using the Orphan-rich dataset as input, predicted more SGD-annotated orphan genes than did either of the *ab initio* predictors or the Direct Inference pipeline alone (**Supplementary Table 4-B**). F1 scores for overall prediction performance were similar for BIND and MIND. Three quarters of the SGD-annotated orphan genes were missed even using the Orphan-rich RNA-Seq datasets as input. However, novel orphan genes were predicted using the Orphan-rich dataset; over 90% of these were predicted by Direct Inference (**Supplementary Figure 3**). BIND and MIND predicted 206 and 223 novel orphan genes, respectively.

### Gene predictions for *Oryza sativa*

The monocot rice is a staple crop for over half the world's population, with a complex genome where 35% of the sequence is made up of transposable elements[3]. We used the MAKER, BRAKER, Direct Inference, BIND, and MIND pipelines to predict rice genes in *Oryza sativa subsp. japonica cv. Kitaake*, and compared these predictions to those of NCBI (GCA_009797565.1). The NCBI annotations are based on the recent high-quality KitaakeX genome[4] and *in silico* gene annotations that were obtained via *ab initio* and homology methods, combined with EST evidence; we were unable to trace which evidence was used for each annotation (Supplementary Table 1). These rice annotations have not undergone the manual community curation as have Arabidopsis and yeast, thus do not provide the same gold standard on which to base our interpretation of the efficacy of the methodologies. *Oryza* phylogenetic research[5,6] has predicted multiple orphan genes; to our knowledge, these predictions have not been integrated into the NCBI annotations. Because the gene annotations we are using for rice have not yet been highly curated, they do not represent the same gold standard as do the Arabidopsis and yeast gene annotations. It is more likely that the annotations include some false positive predictions and are missing some true genes.

We assembled two RNA-Seq datasets to input to the gene prediction pipelines. The first was a "Typical" dataset; the second was a larger "Pooled" dataset consisting of diverse tissues and stress conditions (**Supplementary Tables 2-C and 3-C**). Because the rice gene annotations have not yet been extensively curated manually, we did not assemble an orphan-rich dataset. We partitioned the NCBI-annotated genes, as well as the resultant predictions made by each of the pipelines, according to inferred phylostrata[7] (**Supplementary Tables 7-C and 12**).

Similar to the results with Arabidopsis and yeast, regardless of prediction scenario, the ability to predict genes was greatest for the genes of the most ancient phylostratum and gradually decreased for the younger phylostrata. As observed in Arabidopsis and yeast, more annotated genes were predicted when pipelines were supplied with a more diverse dataset, with the exception being the BRAKER pipeline predictions (**Supplementary Table 4-C**). Similar to Arabidopsis and yeast, MIND predicted more genes matching annotations than did the MAKER or Direct Inference methods alone.

Unlike Arabidopsis and yeast, BRAKER predicted more genes that match to the NCBI annotations compared to any other pipeline (**Supplementary Table 4**). However, 13,178 of the transcripts that BRAKER predicted contained incorrect fusions of splice junctions (**Supplementary Table 9-C**). In contrast, no incorrect fusions were predicted by BRAKER for either Arabidopsis or yeast (**Supplementary Tables 9-A and 9-B**), These 13,178 incorrect fusions were removed by BIND, in the step combining BRAKER and Direct Inference predictions using Mikado. MAKER predicted 2302 incorrect fusions and NCBI annotations include 1374 incorrect fusions. (In contrast, no incorrect fusions were predicted by BRAKER for either Arabidopsis or yeast (**Supplementary Tables 9-A and 9-B**)). About 28% of BRAKER-predicted rice genes, and 13% of BIND-predicted rice genes, are inferred by phylostratal analysis to be orphans; most of these were not annotated by NCBI (**Supplementary Figure 4**).
